# An accelerating, decreasing phylogenetic trend in SARS-CoV-2 genome compositional heterogeneity during the pandemic

**DOI:** 10.1101/2024.12.03.625388

**Authors:** José L. Oliver, Pedro Bernaola-Galván, Pedro Carpena, Francisco Perfectti, Cristina Gómez-Martín, Silvia Castiglione, Pasquale Raia, Miguel Verdú, Andrés Moya

## Abstract

The rapid evolution of SARS-CoV-2 during the pandemic, driven by a plethora of mutations, many of which enable the virus to evade host resistance, has likely altered its genome’s compositional structure (i.e. the arrangement of compositional domains of varying lengths and nucleotide frequencies within the genome). To explore this hypothesis, we summarize the evolutionary effects of these mutations by computing the Sequence Compositional Complexity (SCC) in random datasets of fully sequenced genomes. Phylogenetic ridge regression of SCC against time reveals a striking downward evolutionary trend, as well as an increasing rate of change, suggesting the ongoing adaptation of the virus’s genome structure to the human host. Other genomic features, such as strand asymmetry, the effective number of K-mers, and the depletion of CpG dinucleotides, each linked to the virus’s adaptation to its human host, also exhibit decreasing phylogenetic trends over the course of the pandemic, along with strong phylogenetic correlations to SCC. Overall, our findings suggest an accelerated, genome-wide evolutionary trend toward a more symmetric and homogeneous genome compositional structure in SARS-CoV-2.

## Introduction

Nucleotide frequencies usually vary along the nucleotide chain, resulting in intragenomic biases ^1^. These biases ultimately contribute to the formation of a genome’s compositional structure, which was first uncovered by analytical ultracentrifugation of bulk DNA ^2^, as well as through statistical-physics methods directly analysing long-range correlations in nucleotide sequences (power spectra, fluctuation analysis in DNA walks and entropic sequence segmentation) ^3–5^. The evolution of genome compositional structure has garnered renewed attention in recent years from both theoretical and applied grounds: 1) adequate modelling of compositional heterogeneity is essential for obtaining reliable phylogenetic trees, especially when different lineages exhibit varying nucleotide or amino acid compositions ^6^; 2) considering sequence compositional structure has proven to be highly useful in predicting the emergence of coronavirus Variants of Concerns (VOCs) with enhanced transmission ^7^; and 3) Sequence Compositional Complexity (*SCC*) has, for the first time, enabled the discovery of phylogenetic trends driven by natural selection in Cyanobacteria ^8^.

Compositional heterogeneities range in size from a few nucleotides to tens of millions of them (see ^9,10^ for recent reviews). Arrays of compositional domains of different GC content along the genome sequence form compositional genome structure, which may be associated with important biological features, such as gene and repeat densities, timing of gene expression, or recombination frequency ^2,10^. Genome structure can be changed by any mutational event: point mutations, genome rearrangements, or recombination events. Despite its proofreading mechanism and the brief time-lapse since its appearance, all these changes have been reported in the coronavirus; see ^11^ for recent reviews. Online tracking of SARS-CoV-2 variants and mutations of interest is available on the CoVariants site ^7^ (https://covariants.org/).

Mutational events affecting the structural compositional heterogeneity of a genome can be effectively summarized and quantified by *SCC* ^12^. To achieve this, we first segmented the nucleotide RNA sequence into compositionally homogeneous domains under strict statistical criteria, then accounting for the length and compositional nucleotide differences among the resulting domains by computing its *SCC*. This measure has been recently employed to determine genome complexity in an ancient and diverse group of organisms, the phylum Cyanobacteria ^8^, providing the first evidence for driven evolution towards increasing complexity of genome compositional structure. Tracking changes in sequence compositional structure of coronavirus genomes over time may be relevant on evolutionary and epidemiological grounds. Specifically, the existence of evolutionary trends in the sequence compositional structure of coronavirus genomes could reveal whether natural selection is providing adaptation of the virus’s genome structure to the human host.

In this paper, we computed *SCC* in random datasets of high-quality, completely sequenced coronavirus genomes free of ambiguity symbols (mainly N, R, Y, S, W). Then, we calculated their phylogenetic ridge regression over time ^13^. This method has proven effective in revealing both morphological ^14^ and genomic evolutionary trends ^8^ trends. We present evidence for long-term adaptive tendencies, indicating a decreasing trend in *SCC* and an increasing trend in its evolutionary rate, suggesting that the reduction in *SCC* accelerated over time. Additionally, we sought links between changes in genome compositional structure and other biological features potentially linked to the virus’s adaptation to its human host, such as strand asymmetry, the effective number of *K*-mers, and CpG depletion, which might suggest evolutionary trends toward a more symmetric and homogeneous compositional structure in the SARS-CoV-2 coronavirus genomes during the pandemic.

## Results

### Compositional genome structure of the coronavirus

The presence of a compositional structure in the coronavirus was first suggested based on detrended fluctuation analyses ^15^. Here, using entropic compositional segmentation ^16,17^, we found that the SARS-CoV-2 genome effectively consists of an array of statistically homogeneous compositional domains with varying lengths and nucleotide frequencies. In particular, the reference genome sequence (hCoV-19/Wuhan/WIV04/2019|EPI_ISL_402124|2019-12-30) consists of seven compositional domains, resulting in a *SCC* value of 5.7 x 10E^-3^ bits by sequence position. From then on, descendent isolates presented substantial variation in each domain’s number, length, and nucleotide composition. In the dataset of 4,336 completely sequenced coronavirus genomes analysed here, the number of segments ranges between 6 and 10 (Supplementary Table 1). Note that genomes with seven segments are the most frequent, while those with 6 or 10 segments occur at lower frequencies. On the other hand, *SCC* ranges between 4.9 × 10^-3^ and 9.2 × 10^-3^ bits per sequence position on average. Therefore, coronavirus genomes show sufficient compositional variation, as detected by SCC, to infer their genealogical or evolutionary relationships.

The strain name, collection date, SCCs, number of segments, asymmetry indexes, and other measures for each analysed genome are shown in Supplementary Table 2. The sample includes coronavirus VOCs (Alpha, Delta, and Omicron), minor Variants (Beta, Gamma, Kappa, Iota), as well as no-Variant genomes (Supplementary Table 3). The density of coronavirus VOCs and other genome groups in the sample has changed sequentially throughout the pandemic, with Alpha first appearing in 2020, Delta in 2021, and Omicron dominating from 2022 onward (Supplementary Figure 1). A stacked graphical visualization map of the array of segments obtained from each genome is shown in Supplementary Figure 2. The compositional landscape of the coronavirus genome is dominated by six long segments, with shorter, less visible segments scattered along the sequence. A zoomed-in section of the stacked map highlights the variation in segment boundaries across different genomes. Also note the accumulation of GC-rich segments in the 5’ and 3’ regions of the genome sequence.

### Phylogenetic evolutionary trends

We began investigating evolutionary trends of *SCC* in SARS-CoV-2 early in the pandemic (April 2020). In the first samples retrieved from the Global Initiative on Sharing all Influenza Data (GISAID)^18–20^ we found no statistical support for phylogenetic trends. However, with the emergence of the first Variants in December 2020, the phylogenetic ridge regression slope of *SCC vs*. time started to decrease significantly. Nevertheless, many of those early GISAID entries have ambiguous symbols (mainly N, R, Y, S, and W), which complicated downstream analyses, such as the compositional segmentation of a sequence. To overcome this challenge, we have now chosen to exclusively analyse fully sequenced genomes (i.e., those free of ambiguity symbols). Here, we present results from a random dataset of completely sequenced genomes from around the globe, with collection dates spanning from December 2019 to January 2024. A list of the genome sequences retrieved from the GISAID/Audacity database ^18,19^ was compiled as GISAID EPI_SET_240824vr being available at https://doi.org/10.55876/gis8.240824vr. The obtained *SCC* values, number of segments, collection dates, and other relevant data are shown in Supplementary Table 2.

To infer the phylogeny, coronavirus genome sequences were aligned using *MAFFT* ^21^ (with the options *thread -1* and *nomemsave*). The best ML tree was inferred using *IQ-TREE 2* ^22^ using the GTR nucleotide substitution model ^23,24^ (with the options *GTR+F+R2*). To solve polytomies, we used the function *fix.poly* from the *RRphylo* package ^13^. The least-square dating (LSD2) method ^25^ was used to build a time-scaled tree. Finally, we rooted the inferred time tree to the GISAID coronavirus reference genome (hCoV-19/Wuhan/WIV04/2019|EPI_ISL_402124|2019-12-30). To search for phylogenetic evolutionary trends, we used the function *search.trend* of the RRphylo R package ^13^, contrasting the realized slope of *SCC* versus time regression to a family of 1,000 slopes generated under the Brownian motion (BM) model of evolution, which models evolution with no trend in either the *SCC* or its evolutionary rate. A decreasing phylogenetic trend was detected for *SCC* (Fig. 1, left). We have also investigated the phylogenetic trend of the partial complexity *SCC_RY* (Fig. 1, right), which is one of the partial complexities in which *SCC* can be decomposed ^26^. The behaviour of *SCC_RY* is potentially attractive because it mainly reflects strand asymmetries in the distribution of purine/pyrimidines along the genome sequence, which have been related to key biological mechanisms, including protein binding preferences, transcription factor interactions, retrotransposition, DNA damage and repair preferences, transcription-replication collisions, and mutagenesis mechanisms ^27^.

**Figure 1.**
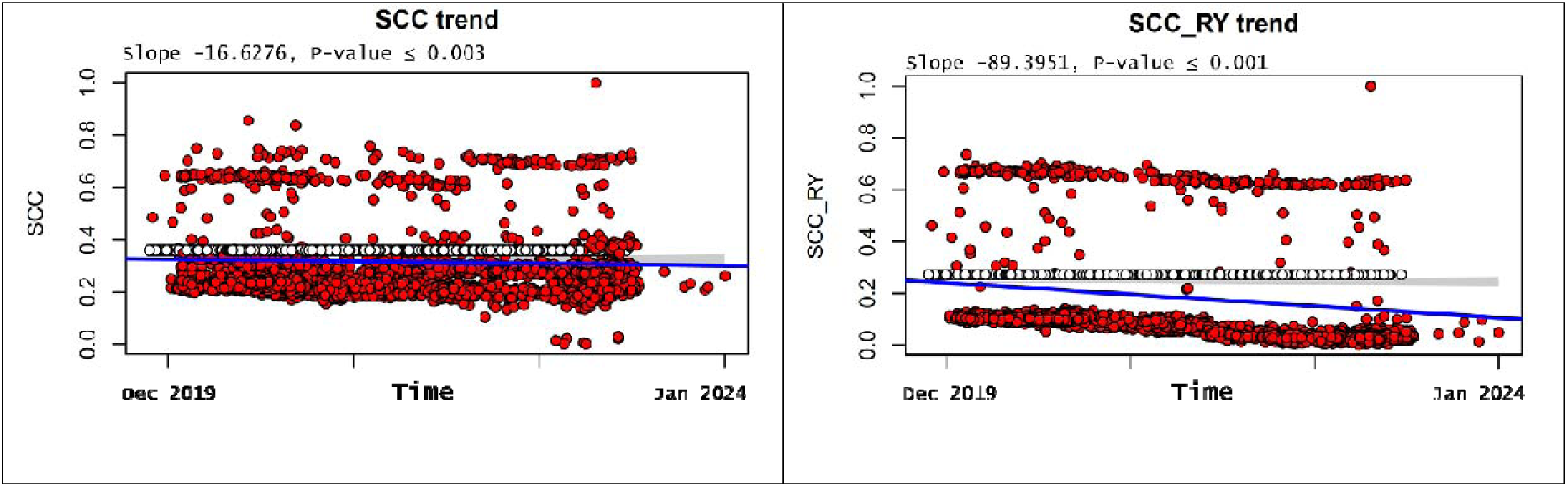
Phylogenetic regressions of *SCC* (left) and the partial complexity *SCC_RY* (right) against pandemic time (days since the first SARS-CoV-2 was isolated), as determined by the RRphylo package ^13^.

The evolutionary rates of *SCC* and *SCC_RY* are shown in Supplementary Figure 3. Both rates increased over time, which means that, alongside the temporal decline in compositional genome complexity, its absolute evolutionary rate increased, suggesting that the reduction in *SCC* and *SCC_RY* accelerated over time. That reinforces the view that the decrease in both *SCCs* (i.e., the homogenization of compositional genome structure) may be part of the virus’s adaptive process to the human host. Finally, note that two compositional groups with low and high compositional complexities are distinguished in the phylogenetic regression plots (Fig. 1) and their respective rates (Supplementary Fig. 3). These compositional differences are likely due to the variation in the number of segments between both groups. A demonstrative example is provided in Supplementary Figure 4 for *SCC_RY*.

### Biases in K-mer distribution

To gain insight into the biological significance of the observed compositional evolutionary trends, we investigated other genomic features that follow similar temporal dynamics. The first of these features is the bias in the distribution of *K*-mers, which are substrings of length *K* that serve as fundamental units for analysing and comparing genomic sequences. The distribution of *K*-mers within a genome sequence holds significant biological relevance, as it provides insights into genomic compositional structure and function ^28,29^.

### Strand asymmetry

To find biases in the distribution of *K*-mers in the coronavirus genome, we first used the S^1^ asymmetry index ^30^ for *K* = 1 to 6. Using ridge regression of S^1^ against time, we observed a highly significant decreasing phylogenetic trend for *K* = 1 (Fig. 2, left) and, to a lesser extent, for *K* = 3 (not shown).

**Figure 2.**
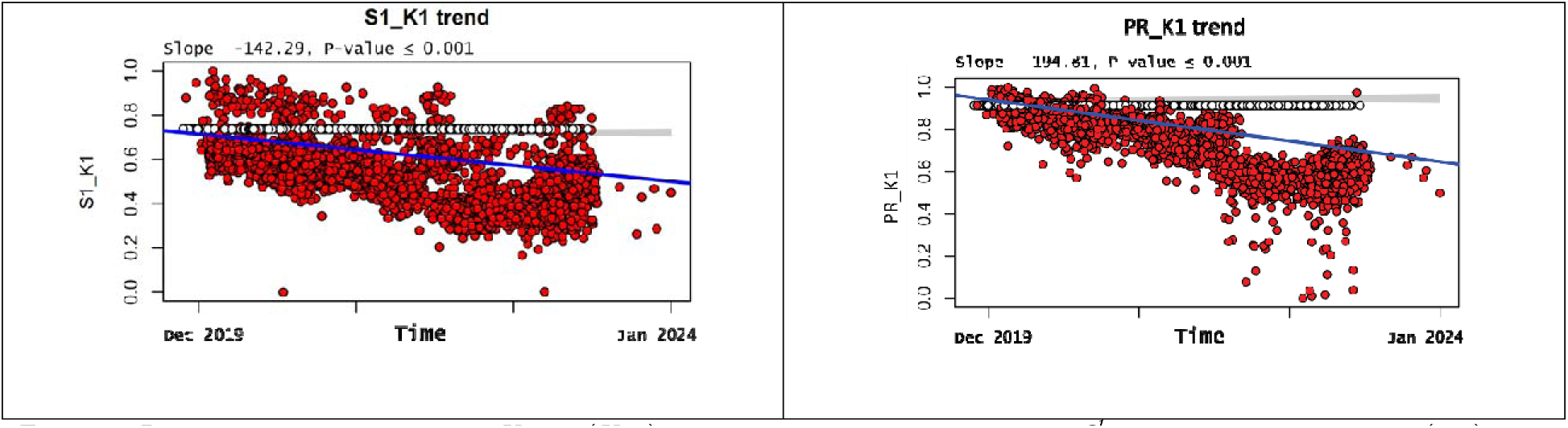
Phylogenetic regression for *K*-mer (*K*=1) distribution, as measured by the *S^1^* strand asymmetry index (left) and the Participation Ratio (*PR*, right), reveals strong phylogenetic decreasing trends over time. The RRphylo package ^13^ was used.

### The Participation Ratio

A second bias in the distribution of K-mers that we used was the effective number of K-mers, computed using the Participation Ratio (PR). We observed highly significant decreasing trends for all *K* values, save *K* = 2. The steepest negative slope was observed for *K* = 1 (Fig. 2, right), which suggests that the number of *K-*mers effectively used by the coronavirus decreased during the pandemic, contributing to the progressive simplification and homogenization of the coronavirus genome.

### CpG depletion

Single-stranded RNA viruses replicating in vertebrate hosts tend to have a low frequency of CpG dinucleotides in their genomes ^31^. Moreover, in SARS-CoV-2, a gradual decline in CpG content has been observed ^32^, albeit at a modest rate over time. Interestingly, when we applied phylogenetic regression to CpG frequencies in our coronavirus dataset, we found a weak but significant decreasing phylogenetic trend (Figure 3, left).

**Figure 3.**
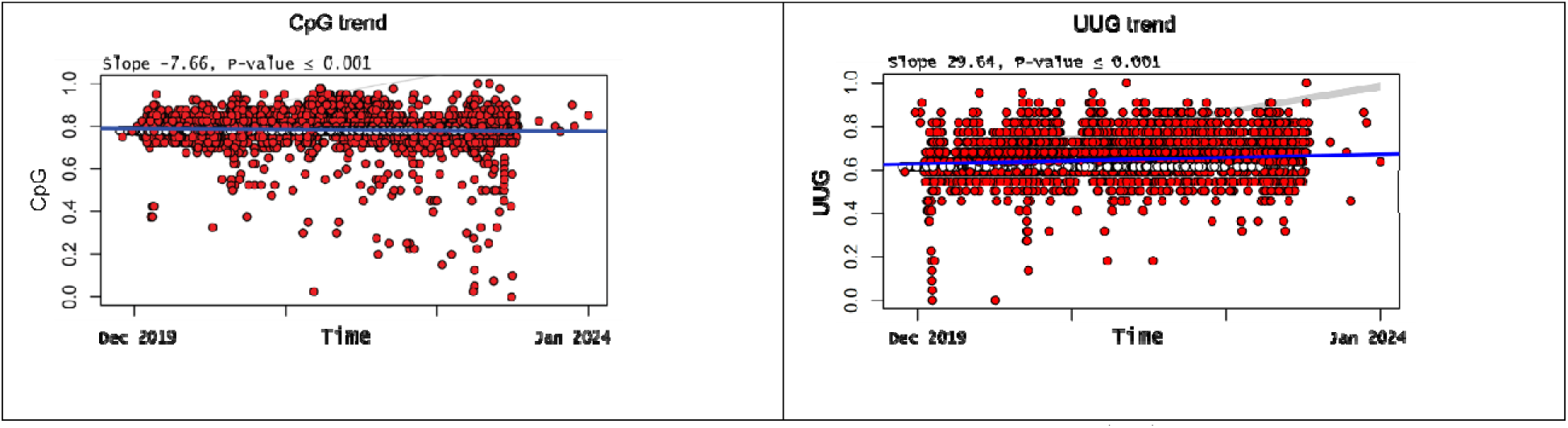
A modest but significant decreasing phylogenetic trend in CpG frequencies (left) and a highly significant increasing trend in the frequencies of their deamination product (UUG) (right) were observed in coronavirus genomes. The RRphylo package ^13^ was used.

Two possible explanations for CpG depletion, based on the directional mutational pressures exerted on the SARS-CoV-2 genome by host antiviral defence systems, have been proposed ^32^. The first attributes CpG depletion to the deamination of methylated cytosines by the host methyltransferases ^33^. However, as SARS-CoV-2 does not have a DNA stage, this mechanism is unlikely to be relevant in the coronavirus ^34^. A second biological mechanism ^35,36^ more clearly explaining the decreasing trends in *SCCs* may be the combined actions of APOBEC (apolipoprotein B mRNA editing catalytic polypeptide-like proteins, which are zinc-dependent deaminases ^37^) and ZAP (zinc-finger antiviral protein ^38^). The catalytic activity of APOBEC enzymes leads to the transformation of 5’-UCG-3’ sites into 5’-UUG-3’ via cytosine deamination, effectively removing the ZAP recognition site (5’-CG-3’). This deamination, changing C!ZU, enables viral RNA to evade degradation by ZAP. Over time, a decrease in CpG dinucleotides and a corresponding increase in their deamination product (UUG) are expected. The decreasing phylogenetic trend we observed in CpG frequencies (Fig. 3, left), coupled with the corresponding increase in UUG trinucleotides (Fig. 3, right), aligns well with this mechanism.

### Phylogenetic correlations of *SCCs* to other biological features

To further investigate the association of *SCCs* to the other biological features with similar temporal dynamics, we constructed a Phylogenetic Generalized Least Squares (PGLS) model ^39^ for each of the four *SCCs* as the dependent variable, and strand asymmetry (*S^1^* index, *K* = 1), the effective number of *K*-mers (*PR, K* = 1) and the frequencies of *CpG* dinucleotides as the independent variables. The R packages *ape* ^40^, *caper* ^41^ and *nlme* ^42^ were used together for this task. The aim here is to understand the significance of each independent on the dependent variable while accounting for phylogenetic relationships. All the phylogenetic correlations obtained from the PGLS models were highly significant (P ≤ 0.001), with the best fit among the four models being obtained for *SCC_RY* (Table 1) since it has the lowest value for the Akaike Information Criterion (*AIC*). This model explains a substantial portion of the variance (81.73%), providing a reliable explanation for the variation in *SCC_RY* across coronavirus genomes. Additionally, the results in Table 1 suggest that *S^1^_K1* has the strongest positive effect, while *PR_K1* shows a negative impact, both being important predictors of *SCC_RY*. The PGLS models for *SCC*, *SCC_SW*, and *SCC_KM* are presented in Supplementary Tables 4–6.

**Table 1.** PGLS model with *SCC_RY* as the dependent variable and *S^1^_K1* (S^1^, *K* = 1), *PR_K1* (PR, *K* = 1), and the frequencies of *CpGs* as the independent ones: PGLS(formula = *SCC_RY* ∼ S^1^_K1 + PR_K1 + CpG). Multiple R-squared: 0.8173, Adjusted R-squared: 0.8172, F-statistic: 6459 on 3 and 4332 DF, P-value: < 2.2e-16.

|  | Estimate | Std. Error | t-value | P-value |
| --- | --- | --- | --- | --- |
| (Intercept) | 3.59E-01 | 1.81E-02 | 19.895 | 0.001 |
| $S^I\_KI$ | 4.88E-01 | 5.00E-03 | 97.727 | 0.001 |
| *PR_KI* | -2.24E-01 | 4.21E-03 | -53.039 | 0.001 |
| *CpG* | 3.71E-05 | 9.70E-07 | 38.259 | 0.001 |

## Discussion

The great number of point mutations, genome rearrangements, and recombination events observed in SARS-CoV-2 have resulted in notable diversification of the virus as it adapted to the human host during the pandemic ^43,44^. Many of these changes, particularly those leading to the emergence of VOCs, may be adaptive. Examples include inter-lineage recombinants ^45^, mutations enabling VOCs to neutralize host resistance or escape antibodies ^46^, consequently increasing transmissibility (a paradigmatic example being the outbreak of the Omicron Variant), co-mutations ^47^ that become more prevalent worldwide compared to single mutations, primarily responsible for temporal changes in transmissibility and virulence, as well as parallel mutations in multiple independent lineages and Variants ^48^, which are of particular interest in the context of adaptation of SARS-CoV-2 to the human host. Structural mutations revealed by homology modelling experiments, which can potentially alter the number or nucleotide frequencies within the array of compositional segments of the coronavirus genome, as well as higher-fitness mutations, such as those in the *nucleocapsid* or *spike* genes, along with hitchhiking mutations in other genomic regions, may also play a role ^49^.

We focus here on the potential effects that all these changes may have had on the evolution of the compositional genome compositional structure of SARS-CoV-2. To this end, we computed *SCC*^12^ and *SCC* partial complexities ^26^, capturing the evolution of the virus’s genome structure in near real-time. Despite its short length (∼29,900 nt), the coronavirus genomes analysed here are segmented into 6 to 10 compositional domains (∼0.25 segments by 1000 nt on average; see the column *nseg* in the Supplementary Table 2). Although such segment density is lower than in free-living organisms (like cyanobacteria, where an average density of 0.47 segments by 1000 nt was observed ^8^), the compositional variability we found in the coronavirus may be sufficient for comparative evolutionary studies of genome structure, which could shed light on the origin and evolution of the COVID-19 pandemic ^49,50^.

Phylogenetic ridge regression of *SCC* and *SCC_RY* over time revealed decreasing evolutionary trends in sequence compositional complexity (Fig. 1), along with increasing rates of change (Supplementary Fig. 3), suggesting the ongoing adaptation of virus’s genome structure to the human host. Notably, applying the same method to other genomic features with similar temporal dynamics— such as strand asymmetry (Fig. 2, left), the effective number of *K*-mers (Fig.2, right), and *CpG* depletion (Fig. 3, left), all of which are potentially linked to key biological features—also reveals decreasing phylogenetic trends over time. The relationships between SCCs and these other biological features were checked by PGLS models^39^, where each *SCC* served as the dependent variable, and strand asymmetry, the effective number of *K*-mers, and *CpG* depletion were the independent variables. The model fit for *SCC_RY* (Table 1) shows the lowest AIC value, indicating that it provides the best explanation for the variation in this partial complexity across the coronavirus phylogeny.

The decreasing phylogenetic evolutionary trends observed in *SCC, SCC_RY*, strand asymmetry, the effective number of *K*-mers, and CpG depletion (Figures 1-3) are particularly interesting, as they suggest that the virus’s ongoing adaptation to the human host has been accompanied by a significant reduction in genome compositional complexity within the global coronavirus population, which points to a progressive simplification and homogenization of the coronavirus genome’s compositional structure over time. Since *SCCs* integrate the complexity of the entire viral genome, reductions in *SCCs* could imply that natural selection is favouring more streamlined, less complex coronavirus genomes over time. This is consistent with the observed decrease in strand asymmetry, which may indicate optimization of replication efficiency across the genome, with selective pressure favouring specific nucleotide compositions to enhance viral fitness^31,51^. Overall, our findings suggest an evolutionary, genome-wide trend toward a more symmetric and homogeneous compositional structure in the SARS-CoV-2 genome, reflecting an adaptive process driven by natural selection as the virus continues to specialize in the human host.

In conclusion, we prove that the increase in fitness of Variant genomes, associated with higher transmissibility, may have contributed to a reduction in SARS-CoV-2 sequence compositional heterogeneity over the course of the pandemic. This genome compositional dynamic may have been driven by the rise of highly fit viral variants and convergent evolution, contributing to an adaptive specialization process in the human host through natural selection. Similar processes have been observed in codon usage and amino acid preferences in other viruses ^52^. Further monitoring of these evolutionary trends in current and emerging Variants and recombinant lineages ^53,54^, using the methodology applied here may help clarify whether—and to what extent—the evolution of compositional genome structure in this and other pathogen genomes affects human health.

## Methods

### The genome of SARS-CoV-2 coronavirus

The SARS-CoV-2 genome is an approximately 30 kb, positive sense, 5’ capped single-stranded RNA molecule ^55^. An updated genomic map of the isolate Wuhan-Hu-1 (MN908947.3) of SARS-CoV-2 we used as a reference genome for alignment is available at https://www.ncbi.nlm.nih.gov/nuccore/MN908947.3?report=graph. Genomic information on the official reference sequence employed by GISAID (EPI_ISL_402124, hCoV-19/Wuhan/WIV04/2019, (WIV04)) is available at https://www.gisaid.org/resources/hcov-19-reference-sequence/. We used this genome as the root when inferring a phylogeny. Note that although WIV04 is twelve nucleotides shorter than Wuhan-Hu-1 at the 3’ end, the two sequences are identical in practical terms; mainly, the 5’ UTR is the same length, and the coding regions are identical. Therefore, the coordinates and relative changes stay the same whichever sequence is used, which is also relevant for coordinates of compositional segments.

### Retrieving a coronavirus dataset free of Ns and other ambiguous symbols

Genome coronavirus sequences are available from GISAID, but many of them are not fully sequenced and have ambiguous symbols (N, R, Y, S, W, K, M), which could complicate the compositional segmentation of a sequence. To overcome this difficulty, we first retrieved a random sample of 10,000 high-quality genomes from around the globe from this database and then discarded duplicates and entries with incomplete collection dates. By using *seqtk* (https://github.com/lh3/seqtk) and *Nextclade* ^56^ computer programs, we further filtered this sample by discarding sequences containing Ns and other ambiguities, resulting in a final dataset of 4,336 unique entries, completely sequenced, free of ambiguous symbols, and spanning from December 2019 to January 2024. This retrieval workflow was used to obtain various datasets, ensuring the repeatability of our analyses.

### Sequence Compositional Complexity (*SCC*)

The sequence compositional structure of each coronavirus genome was determined by computing its Sequence Compositional Complexity (*SCC*) ^12^, which consists of a two-step process: the nucleotide sequence was first segmented into homogeneous, statistically significant compositional domains, followed by the computation of *SCC*. Using the alphabet {A, T, C, G} (remember that in RNA genomes, the letter T is used to denote Uracil (U)), we divided each coronavirus sequence into an array of compositionally homogeneous, non-overlapping domains using a heuristic, iterative segmentation algorithm ^16,17^. In brief, a sliding cursor is moved along the sequence, and the position that optimizes a proper measure of compositional divergence between the left and right parts is selected. We choose the Jensen-Shannon divergence (equations [1] and [2] in ^16^) as the divergence measure, as it can be directly applied to symbolic nucleotide sequences. If the divergence is statistically significant (at a given significance level that we choose to be *s =* 0.95), the sequence is split into two segments. Note that each pair of resulting segments is more homogeneous than the original sequence. The two new segments are then independently subjected to another round of segmentation. The process continues iteratively over the new segments while sufficient significance continues appearing.

Note that the *s* value (here 0.95) is the probability that the difference between adjacent domains is not due to statistical fluctuations. Recent improvements to the segmentation algorithm ^57^ allow segmenting sequences with long-range correlations, as those recently reported in the coronavirus ^15^.

The result is the segmentation of the original sequence into an array of contiguous, non-overlapping segments (or compositional domains) whose nucleotide composition is entropically homogeneous within a predefined level of statistical significance, *s.* A stacked map of the segmentations observed in all the genomes within the analysed dataset is presented in Supplementary Figure 2.

Once a sequence is segmented into an array of homogeneous compositional domains at a given significance level (e.g., P-value ≤ 0.05), a measure of Sequence Compositional Complexity or *SCC* ^12^, expressed in bits by sequence position, was computed:

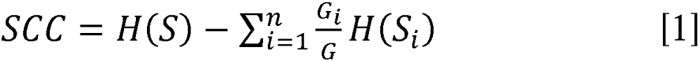

where *S* denotes the whole genome sequence, G is its length, and G_i_ is the length of the *i*^th^ domain 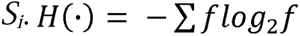 is the Shannon entropy of the distribution of relative frequencies of symbol occurrences, *f*, in the corresponding (sub)sequence. It should be noted that the above expression is the same as the one used in the segmentation process, applying it to the tentative two new subsequences (*n* = 2) to be obtained in each step. In this way, the segmentation procedure finds the partition of the sequence that maximizes *SCC*. It is also worth noting that the two steps of the *SCC* computation are based on the same theoretical background. Note that 1) this measure is zero if no segments are found in the sequence (the sequence is compositionally homogeneous, e.g., a genuinely random sequence) and 2) it increases/decreases with both the number of segments and the degree of compositional differences among them. In this way, the *SCC* measure is analogous to the method proposed by ^58^ for estimating complexity in morphological characters: an organism is more complex if it has a greater number of parts and a higher differentiation among these parts. It is important to emphasize the high sensitivity of this measure to sequence changes. A single nucleotide substitution or one little indel could potentially alter the number, length, or nucleotide frequencies of the compositional domains and, therefore, the resulting value for *SCC*. A Python script to segment the coronavirus genome sequences and compute *SCC* is available at the open repository Zenodo.

### *SCC* partial complexities

The quaternary alphabet {A, T, C, G} is commonly used for *SCC* computation. However, taking advantage of the branching property of entropy ^59^, SCC can be decomposed into partial complexities by grouping the nucleotides into binary alphabets, as *SW*{GC/AT}, *RY*{AG/TC} or *KM*{AC/TG} ^26^. Two of the partial complexities obtained in this way (*SCC_SW* and *SCC_RY*) have been directly associated with key biological features. *SCC_SW* directly reflects changes in GC content, which are often associated with gene and repeat densities, timing of gene expression, or recombination frequency ^2,10^ *SCC_RY* mainly reflects strand asymmetries in the distribution of purine/pyrimidines along the sequence, being related to key biological mechanisms, including protein binding preferences, transcription factor interactions, retrotransposition, DNA damage and repair preferences, transcription-replication collisions, and mutagenesis mechanisms ^27^. Nonrelevant biological features have been associated with the alphabet KM {AC/TG} ^60^.

### Phylogenetic ridge regression

The phylogenetic ridge regression of *SCC* was determined by using the *RRphylo* R package ^13^. In *Rrphylo,* the change in *SCC* value between any two consecutive tree branches aligned along a phyletic line is described by the equation *ΔSCC = β_1_1_1 +_ β_2_1_2 +_ … _+_ β_n_1_n_* where the *β_ith_* and *l_ith_* elements represent the regression coefficient and branch length, respectively, for each *i_th_* branch along the phyletic line. The matrix solution to find the vector of *β* coefficients for all the branches is given by the equation 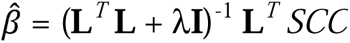; where **L** is the matrix of tip-to-root distances of the tree (the branch lengths), having tips as rows, where entries are zeroes for the branches outside the tip phyletic line, and actual branch lengths for those branches along the path. *λ* is a penalization factor that avoids perfect predictions of *SCC,* preventing model overfitting. The vector of ancestral states 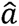 (*SCC* values at the tree nodes) is obtained by the equation 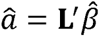 , where L^’^ is the node-to-root path matrix, calculated as **L**, but with nodes as rows. The estimated *SCC* value for each tip or node in the phylogenetic tree is regressed against its age (the phylogenetic time distance, which represents the time distance between the reference genome and the collection date of individual virus isolates) and the regression slope compared to Brownian Motion (*BM*) expectations (which predicts no trend in *SCC* values and rates over time) by generating 1,000 slopes simulating BM evolution on the phylogenetic tree, using the function *search.trend* ^61^ in the *Rrphylo* R package.

### Measuring strand asymmetry (S^1^)

Strand asymmetry for each coronavirus genome sequence was computed using its distribution of *K*-mers. One popular method ^30^ first computes the *K*th-order strand symmetry of any given sequence as the similarity between its *K*-mer distribution f and the *K*-mer distribution *f’* of its actual or virtual reverse complement. Let us consider the standard four-letter alphabet {A, T, C, G}, then there are 4*^K^*different *K*-mers. Given the observed distribution of *K*-mers in the analysed sequence, if *f*_i_ stands for the relative frequency of the i-th *K*-mer, then 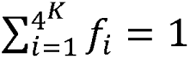. In practice, this method used the sum of the absolute values of the differences between *K*-mer frequencies:

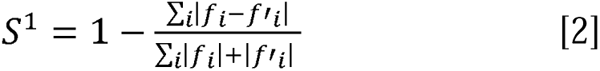

*S*^1^ ranges from 0 (asymmetry/dissimilarity) to 1 (perfect symmetry/similarity). When computed on distributions, it stands for the percentage of *K*-mer occurrences that are symmetrically distributed among complementary strands. Its complement to 1 (an asymmetry index) indeed corresponds to the weighted average of the absolute values of the skews of reverse-complementary bases or *K*-mers.

Baisnée *et al.* ^30^ also propose computing strand symmetry using Pearson’s linear correlation coefficient, *S^C^*, which ranges from -1 to 1 and generally yields results that are qualitatively similar to those obtained with *S^1^*.

### Measuring the Participation Ratio (*PR*)

In genomic sequences, it is widely recognized that over-represented *K*-mers, like stretches of As or Ts (poly(A) and poly(T) tracts), can skew the *S^C^* symmetry index ^30^. Sequences with a more diverse *K*-mer distribution tend to produce higher *S^C^* values. To prevent this bias, we propose another measure able to capture the main characteristics of the *K-*mer distribution: the Participation Ratio, or *PR*. Given an observed distribution of *K*-mers with relative frequencies *f*_i_, the PR for such distribution is calculated as:

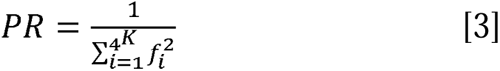

The meaning of *PR* can be understood by considering two extreme situations: If all the *K*-mers appear with the same frequency f = 1/4^K^ then *PR* =4*^K^*, i.e., all *K*-mers contribute (or *participate*) equally to the *K*-mer distribution. If only a single *K*-mer appears in the distribution, then f_i_ =1 for such *K*-mer and f_i_ =0 for the rest, and therefore *PR* = 1 since only one *K*-mer participates in the distribution. In general, *PR* indicates the number of *K*-mers participating effectively in the observed distribution. *PR* is commonly used in quantum solid-state physics to calculate the number of atoms where an electronic wave function is markedly different from 0 (see, for example, ^62^).

## Acknowledgements

We gratefully acknowledge all data contributors, i.e., the Authors and their Originating laboratories responsible for obtaining the specimens and their Submitting laboratories for generating the genetic sequence and metadata and sharing via the GISAID Initiative, on which this research is based. A complete list of the IDs of genome sequences acknowledging all originating and submitting laboratories is available from the GISAID/Audacity database ^18,19^, being available at https://doi.org/10.55876/gis8.240824vr. This project was funded by grants from the Spanish Minister of Science, Innovation and Universities (former Spanish Minister of Economy and Competitiveness) to J.L.O. (Project AGL2017-88702-C2-2-R), P.C. and P.B.G. (Ministerio de Ciencia e Innovación/Agencia Estatal de Investigación, Grant. No. PID2020-116711GB-I00), A.M. (Project PID2019-105969GB-I00), and a grant from Generalitat Valenciana to A.M. (Project Prometeo/2018/A/133) and co-financed by the European Regional Development Fund (ERDF). The most time-demanding computations were done on Linux servers in 1) the Laboratory of Bioinformatics, Dept. of Genetics & Institute of Biotechnology, Center of Biomedical Research, 18100, Granada, Spain; and 2) the Dept. of Applied Physics II and Institute Carlos I for Theoretical and Computational Physics, University of Málaga, 29071, Málaga, Spain.

## Author contributions

J.L.O., M.V., and A.M. designed research; J.L.O., P.B., P.C., F.P., C.G.M, S.C., P.R., M.V. and A.M. performed research. J.L.O., P.B., P.C., F.P., C.G.M, S.C., P.R., M.V., and A.M. analysed data; J.L.O., P.B., P.C., M.V., A.M., and P.R. drafted the paper. All authors have read and approved the final manuscript.

## Data availability

A list of the 4,336 genome sequences analysed here, retrieved from the GISAID/Audacity database:

- GISAID EPI_SET_240824vr available at https://doi.org/10.55876/gis8.240824vr.

The following additional data and scripts are available at the open repository Zenodo (https://doi.org/10.5281/zenodo.13753013):

- A file showing the number and nucleotide composition of the compositional segments in each genome sequence: *segments_4336.txt*
- The rooted timetree in Newick format: *timetree_4336.nwk*
- The Python script used to segment the coronavirus genome sequences and compute SCCs: *scc.zip*
- Script help file: *scc_readme.rtf*

## Additional Information (including a Competing Interests Statement)

The authors declare no competing interests.

