## Supplementary Figures for "An accelerating, decreasing phylogenetic trend in SARS-CoV-2 genome compositional heterogeneity during the pandemic": SupplementaryFigures.docx

### An accelerated, decreasing evolutionary trend in nucleotide compositional heterogeneity of the SARS-CoV-2 genome during the pandemic

### Supplementary figures

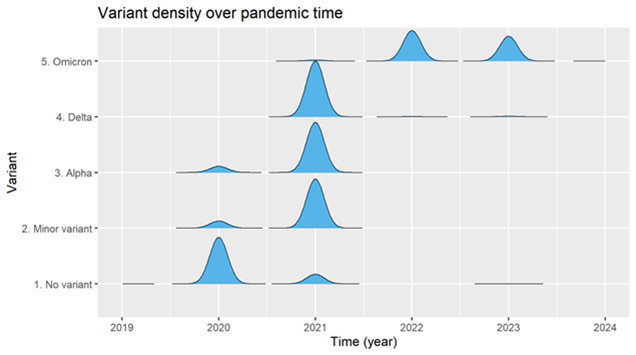

Supplementary Figure 1. Changing densities over time of main VOCs and other genome groups in the sample studied.

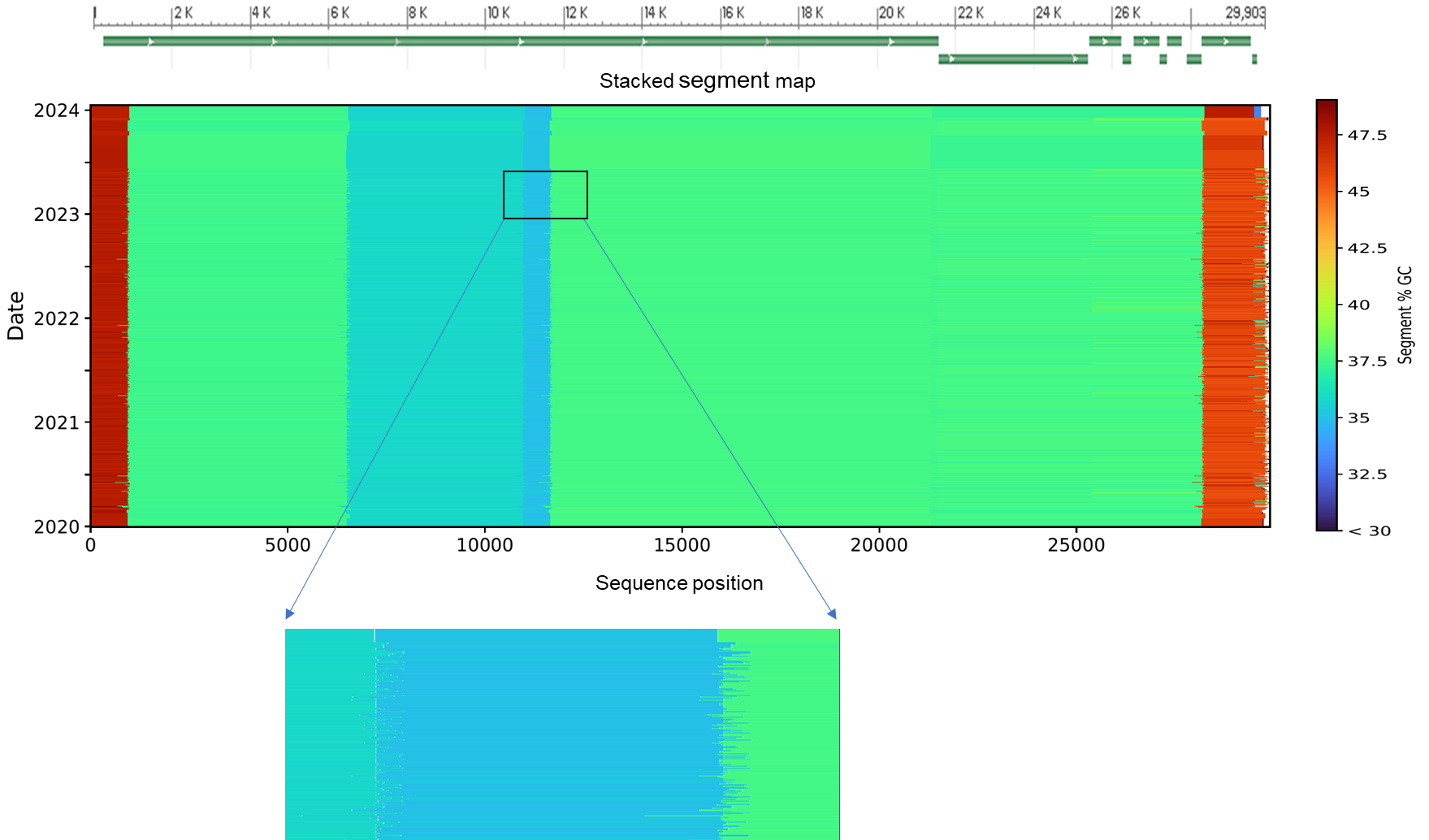

Supplementary Figure 2. Stacked graphical visualization map of the array of segments obtained from each genome, ordered by collection date. Each genome’s array of segments is depicted as a thin horizontal line, with colours standing for the %GC content of each segment (scale on the right). The zoomed region highlights the variation at segment boundaries across different genomes. A schematic gene map of the GenBank reference genome (MN908947.3, Wuhan-Hu-1) is displayed above. A more detailed view of the gene map is available at <https://www.ncbi.nlm.nih.gov/nuccore/MN908947.3?report=graph>.

| 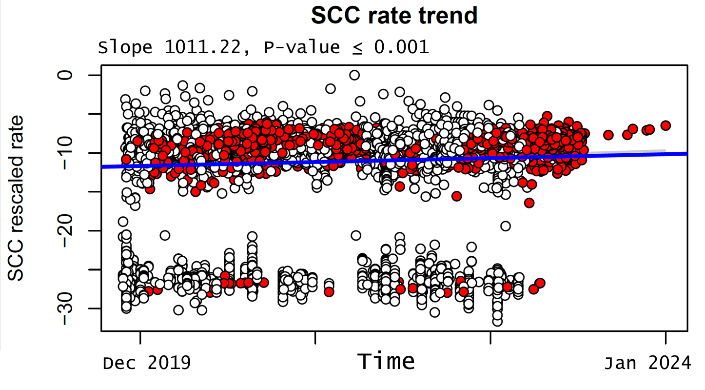 | 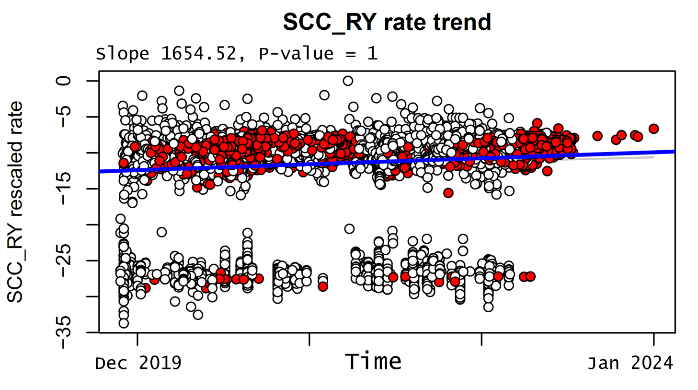 |
| --- | --- |

Supplementary Figure 3. Evolutionary rates of *SCC* (left) and *SCC_RY* (right) over pandemic time.

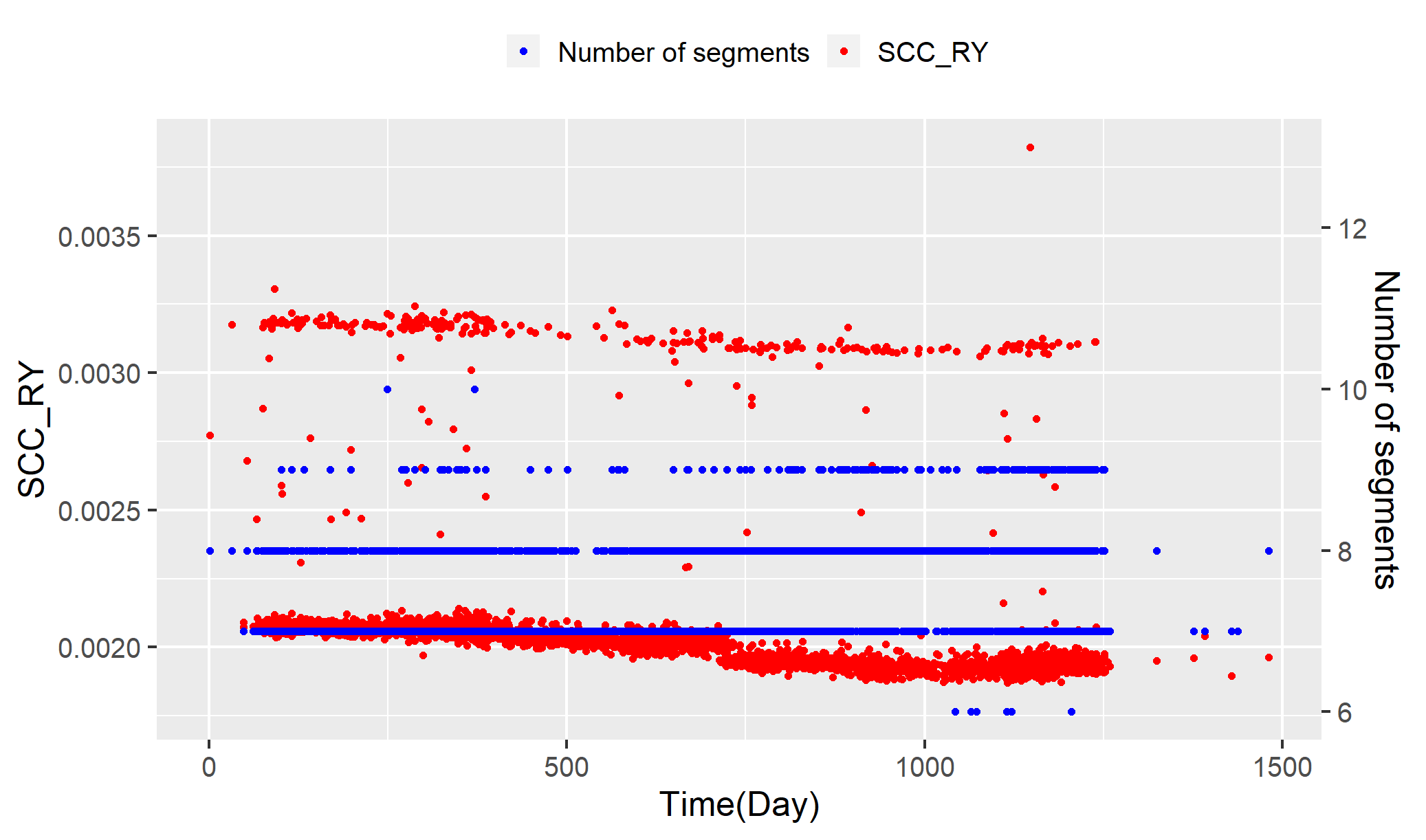

Supplementary Figure 4. The two compositional groups with high or low *SCC_RY* (left Y axis) mainly differ in the number of segments (right Y axis).
